# CancerSiamese: one-shot learning for predicting primary and metastatic tumor types unseen during model training

**DOI:** 10.1101/2020.09.07.286583

**Authors:** Milad Mostav, Yu-Chiao Chiu, Yidong Chen, Yufei Huang

## Abstract

We consider cancer classification based on one single gene expression profile. We proposed CancerSiamese, a new one-shot learning model, to predict the cancer type of a query primary or metastatic tumor sample based on a support set that contains only one known sample for each cancer type. CancerSiamese receives pairs of gene expression profiles and learns a representation of similar or dissimilar cancer types through two parallel Convolutional Neural Networks joined by a similarity function. We trained CancerSiamese for both primary and metastatic cancer type predictions using samples from TCGA and MET500. Test results for different *N*-way predictions yielded an average accuracy improvement of 8% and 4% over the benchmark 1-Nearest Neighbor (1-NN) classifier for primary and metastatic tumors, respectively. Moreover, we applied the guided gradient saliency map and feature selection to CancerSiamese to identify and analyze the marker-gene candidates for primary and metastatic cancers. Our work demonstrated, for the first time, the feasibility of applying one-shot learning for expression-based cancer type prediction when gene expression data of cancer types are limited and could inspire new and ingenious applications of one-shot and few-shot learning solutions for improving cancer diagnosis, treatment planning, and our understanding of cancer.

## 1 INTRODUCTION

More than 200 different cancer types are discovered to originate from different organs and sub-tissues. Yet, molecular signatures of the same cancer type can vary with its location, stage, and ultimately patients. To gain insights into the genetic markers and molecular mechanisms of different cancers, the comprehensive genomic studies such as the Cancer Genome Atlas (TCGA) [1, 2] have generated and interrogated the genetic and omics (epigenomic, transcriptomic, and proteomic) profiles from large cohorts of cancer patients with some of the most common cancer types. It becomes increasingly clear now that the spectrum of cancer transcends existing tumor lineages, underscoring the need for a molecular-based classification of individual tumors. This emergent perspective of cancer fosters the “precision cancer therapy”, which advocates specialized diagnosis and treatments based on the molecular makeup of individual patients [3].

In this paper, we consider cancer classification based on one gene expression profile of known cancer types using machine learning. Thanks to efforts like TCGA, the prevailing strategy is to train a classifier including notably deep learning (DL) using tumor samples with annotated cancer types. To use image-based Convolutional Neural Networks (CNNs), many approaches [4, 5] converted 1-dimensional (1-D) gene expression into 2-dimensional (2-D) image-like inputs or incorporated gene correlations into the converted 2-D images [6–8] before feeding the expression into CNN to classify cancer types. In [9], a 1D-CNN model applied directly to gene expression was tested for classifying 33 TCGA cancer types. It showed that the 1D-CNN could achieve comparable performance with CNNs using 2D-converted gene expression and had 100 times fewer parameters. As the training sample size is commonly small, this simpler 1D-CNN is favored because it has less tendency to overfitting and therefore is more robustness with a better ability to generalize.

However, this strategy has limitations, restricting its adoption in the era of precision oncology [10]. First, the high accuracy is predicated on the availability of large-scale well-annotated tumor datasets like TCGA. Second, for rare tumors, one could never expect to collect sufficient samples necessary for training an ML model with satisfactory performance. Above all, as cancer classification quickly shifts to a more refined molecular-based characterization, we expect to see a much larger number of cancer types. The current strategy is inept with this multitude because every time there is a change in cancer types, the classifier needs to be completely re-trained.

We propose a different one-shot learning approach in this paper, where we only require a single “support” sample from each new cancer type, a drastically reduced requirement from typical TCGA collection with ~500 samples per cancer type. Cancer classification of a query sample is carried out by comparing it against a set of support samples, one for each cancer type (Supplementary **Figure S1**). One-shot learning was first proposed in computer vision for tackling data scarcity [11, 12]. A popular one-shot learning model is Siamese CNN (SCNN), which has also been applied for bioinformatics and medical imaging applications [13–18]. With the success of these studies, we hypothesize that there is a set of type-agnostic marker genes whose expression profiles define the similarity/dissimilarity between samples of the same/different types. Therefore, we shift our attention from predicting the cancer type of the query sample to predicting similarity vs. dissimilarity between a pair of query and support samples. This new perspective enables us to train SCNN with paired samples from the same or different types as replicates for label “similar” or “dissimilar”, thus significantly reducing the need for collecting large samples for each type. Instead, it advocates sampling more types as opposed to sampling more tumors of the same type, a new practice aligned with the nature of precision oncology. Lastly, because the maker genes are type-agnostic, the trained model can be directly applied to classify new cancer types unseen in the training data.

To test this hypothesis, we developed the CancerSiamese, a one-shot SCNN model that contains two identical 1D-CNNs to learn cancer type representations of query and support samples, followed by a metric-learning layer to predict if the query and support sample are similar. We trained and tested CancerSiamese on samples from 29 primary and 20 metastatic cancer types to predict unseen primary and/or metastatic tumor classes and conducted comprehensive investigations of the marker genes learned by CancerSiamese. Our work is noticeably different from AffinityNet [19], a recently developed kNN-based graph attention few-shot learning model, which was also applied for cancer type prediction. AffinityNet is not a one-shot learning model and thus needs more than a single sample for prediction. Also, AffinityNet is very limited in its scope for cancer type predictions. First, it was trained for only two primary cancer types. Second, it was trained to predict only the cancer types that exist in training, so it could not be used to predict novel cancer types. Finally, AffinityNet did not attempt to interpret its predictions and thus did not inform markers and functions underlying the prediction.

## 2 Materials and Method

### 2.1 Dataset

RNA-Seq data from TCGA and Integrative Clinical Genomics of Metastatic Cancer known as MET500 [2] were downloaded. The gene expressions were transformed by log2(FPKM + 1) and filtered based on their mean and variance. We used the tissue of origin to label both primary and metastatic tumors and retained only the common genes between TCGA and MET500. After the preprocessing, we obtained samples from 29 types, where 9 are unique to TCGA (primary) cancers and 20 are common between TCGA and MET500 (primary and metastatic) cancer types (**Figure 1**), where each sample contained 4,858 common genes. The preprocessing is detailed in the supplementary file.

**Figure 1.**
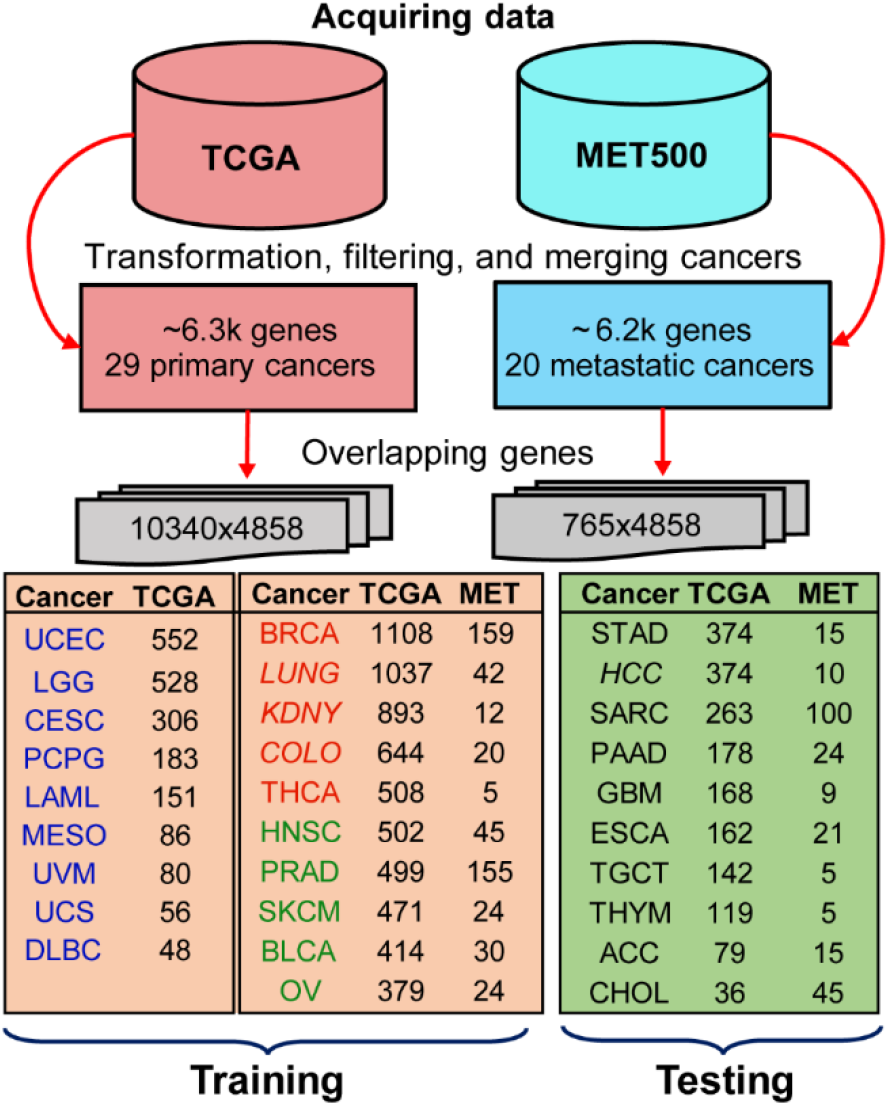
Preprocessing workflow for extracting the training and testing primary and metastatic tumor samples. After downloading the TCGA and MET500 datasets, data preprocessing, including filtering of genes and merging related cancer types were performed. 4,858 common genes in TCGA and MET500 were retained and then samples were divided into training and testing sets according to the number of primary cancer types in TCGA. Cancer types in italic font are merged tumor groups as described in the Methods section. Three training datasets for primary cancers were created and include 9 (blue labeled), 14 (blue + red labeled), and all 19 cancer types, respectively.

The training data include samples from 9 unique primary cancer types and 10 additional cancer types with both primary and metastatic samples (**Figure 1**). The testing data for primary and metastatic cancers include samples from the remaining 10 cancer types (**Figure 1**, green color). It is important to note that the testing set does not share any common cancer types with the training set.

### 2.2 Proposed network model for CancerSiamese

We consider a scenario where we have a query and *N* support gene expression samples, each from a different known cancer type. Among the *N* support samples, one has the same cancer type as the query sample. The goal is to determine which one of *N* cancer types that the query sample belongs to, or make an *N*-way prediction. To achieve this goal, we propose CancerSiamese, inspired by the Siamese network [20]. CancerSiamese takes a pair of query and support samples and then computes the probability that the query is from the same cancer type as the support. After CancerSiamese is applied to all the supporst samples, the query cancer type is predicted as the one with the highest probability (Supplementary **Figure S1**).

CancerSiamese consists of two identical 1D-CNN [9] applied to the query and support sample individually, followed by a similarity metric network (**Figure 2**A). The 1D-CNN includes three 1D convolutional layers and maxpooling layer, and finally a flatten layer. Similar to [20], the relu activation function was selected for the first two 1D convolution layers and sigmoid for the last 1D convolution layer. For the similarity metric network, an elementwise *L*_2_ distance was applied to the two feature vectors generated by the 1D-CNNs and the output was passed onto two consecutive fully connected (FC) layers followed by a sigmoid node to output the probability of similar/dissimilar.

**Figure 2.**
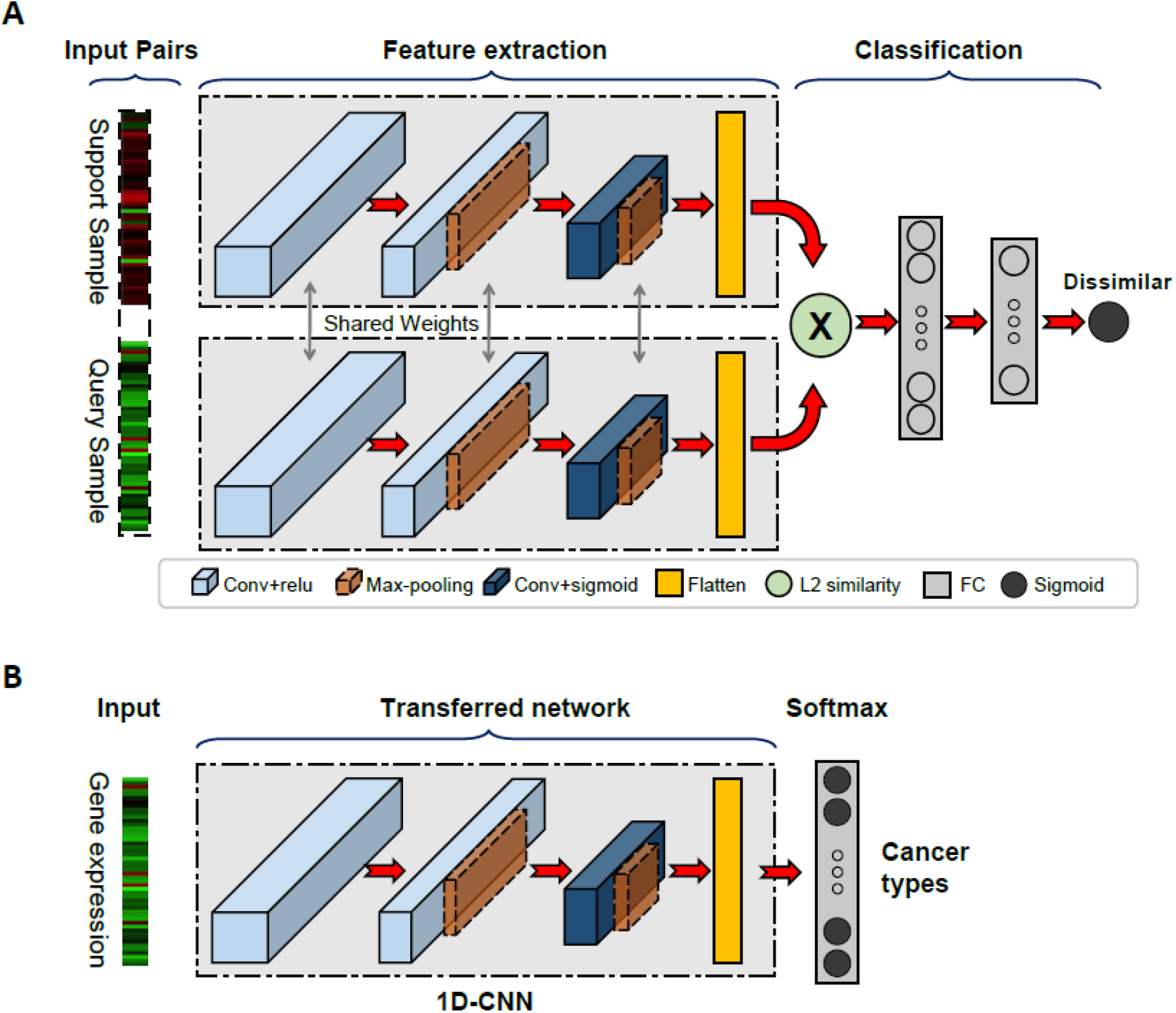
Architecture and network transfer training scheme for CancerSiamese. A) CancerSiamese model architecture. B) The architecture of 1D-CNN, which is pre-trained for cancer classification to initialize the feature extractor part of CancerSiamese.

### 2.3 Network transfer learning

Due to the large network size of CancerSiamese, training the network from scratch suffered from training instability and poor convergence, eventually resulting in less robust prediction. To address these training challenges, we adopted a transfer learning scheme to build the CancerSiamese model. Specifically, we first trained a 1D-CNN using the cancer training samples for cancer type classification. Afterward, we removed the classification layer and took the remaining trained 1D-CNN for CancerSiamese (**Figure 2**B). During the training of CancerSiamese model, these 1D-CNN networks were further optimized for predicting the similarity of the input samples. Because the 1D-CNN has already been trained to extract expression representations important for cancer type classification, the weights are closer to the one-shot learning optimal. Therefore, initializing from this pretrained 1D-CNN makes CancerSiamese training more stable and converging faster.

### 2.4 *Computing the scores of marker genes from* CancerSiamese

We hypothesized that the CancerSiamese trained on primary and metastatic tumors (the details are described in the Results section) rely on maker genes to define a similar cancer type. To uncover the marker genes, a deep learning interpretation approach known as Guided Backpropagation Saliency Maps (GBSM) [21] was employed to compute a score for each gene to be a marker gene by backpropagating a gradient from the output of the model to the input to examine the impact on model’s final decisions. We have previously applied this approach to extract significant cancer markers across different primary cancer types [9]. Briefly, for an input pair of expression samples ***x*_0_** and ***x*_1_**, from the same cancer type, GBSM computed the corresponding gradient vectors ***w*_0_** and ***w*_1_** of the same dimension as the input ***x*** and whose element represents the significance of the corresponding gene. We calculated position-wise average ***w_avg_*** = (***w*_0_** + ***w*_1_**)/2 to obtain a single vector for every pair of expressions from the same cancer type and then computed 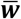 averaged ***w_avg_*** over all the similar pairs used for identifying marker genes, as the scores for each gene to be maker genes.

## 3. RESULT

### 3.1 Model training

CancerSiamese network shown in **Figure 2**A was trained on TCGA and MET500 training datasets separately with Keras DL platform with the Tensorflow backend [22]. To assess the relationship between the number of training cancer types and prediction performance, three CancerSiamese networks for primary cancer prediction were trained using three different sets of primary cancer types, where the total number of cancer types was 9,14, or 19, respectively (i.e., CancerSiamese-9, −14, and −19; **Figure 2**). In contrast, only one CancerSiamese network (CancerSiamese-MET) was trained for metastatic tumor prediction, using samples from 10 metastatic cancer types. Network transfer learning was conducted from the training of each CancerSiamese model, where the initial weights of the 1D-CNN feature extractors (**Figure 3**A) were set as those in the 1D-CNN for classification of cancer types pretrained on the same training set (**Figure 3**B). For example, CancerSiamese-19 took the weights of the 1D-CNN classifier for classifying the same 19 primary cancer classes (**Figure 3**B). The weights for the rest of the layers (i.e. FC and sigmoid) were initialized by Xavier Initialization as suggested by [20]. Each CancerSiamese network was optimized with a binary cross-entropy loss and trained with 20,000 training iterations, where each iteration includes a batch of 128 pairs with an equal number of matched and mismatched pairs, all chosen randomly from the corresponding training dataset (**Table 1**). The network parameters were optimized by Adam optimizer and all of the hyperparameters were tuned manually and summarized in Supplementary **Table 2** and **Table 3**.

**Figure 3:**
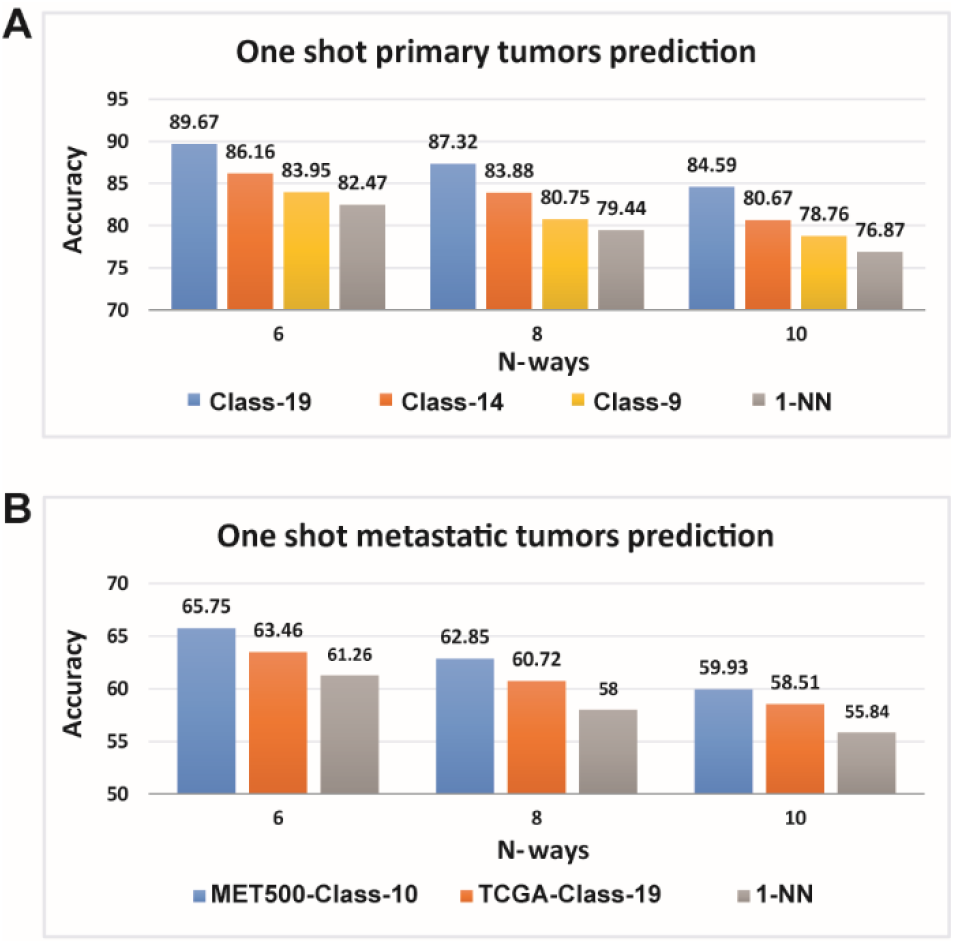
Performance of CancerSiamese for predicting unseen primary and metastatic cancer types. A) Accuracies for predicting primary tumor types. B) Accuracies for predicting metastatic tumor types. C) Confusion matrix of 1D-CNN for cancer type classification trained on 10 primary cancer types and tested on corresponding 10 metastatic tumors in the training set.

**Table 1:**
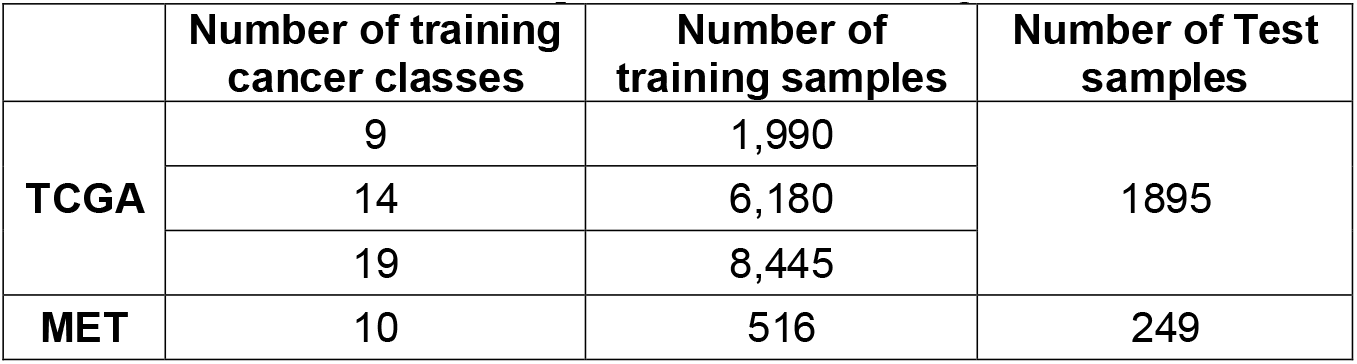
Number of samples for different training and test datasets

|  | Number of training cancer classes | Number of training samples | Number of Test samples |
| --- | --- | --- | --- |
| **TCGA** | 9 | 1,990 | 1895 |
|  | 14 | 6,180 |  |
|  | 19 | 8,445 |  |
| **MET** | 10 | 516 | 249 |

**Table 2:** Accuracies of the SGFS for selecting marker genes for primary cancers (column 2) and their ability for predicting metastatic cancer types (column 3) and discriminating primary from metastatic cancers (column 4)

| Number of top genes | 6-way prediction (Primary cancers; 1-NN) | 6-way prediction (Metastatic cancers; 1-NN) | Classification of primary and metastatic cancers (GNB) |
| --- | --- | --- | --- |
| 10 | 78.64 | 37.4 | 88.35 |
| 20 | 80.74 | 38.11 | 95.22 |
| 30 | 81.61 | 37.58 | 92.87 |
| 40 | 81.8 | 38.12 | 93.13 |
| 50 | 81.8 | 37.29 | 93.55 |
| 100 | 82.38 | 37.51 | 93.28 |
| 150 | 82.23 | 37.08 | 94.38 |
| 200 | 82.1 | 37.36 | 95.1 |

The trained CancerSiamese networks were tested for different *N*-way predictions (*N* = 6, 8, and 10). For an *N*-way prediction, CancerSiamese compared a query sample with a support set of *N* samples, each from a different cancer type. The type of the query sample was predicted as the type of the paired support sample if the pair received the highest probability by CancerSiamese out of the *N* pairs. The prediction was counted as correct if the predicted type was the same as the true type of the query sample. For every *N*-way prediction, we tested CancerSiamese on 20,000 randomly selected query samples and the corresponding support set from the test dataset. Each support set contained *N* randomly selected samples, each from a different cancer type but one of them coming from the same cancer type as the query sample. Finally, the accuracy of the model was calculated to measure the performance. The total number of samples and classes for each training and test datasets are presented in **Table 1**. All of the codes can be found at

### 3.2 Predicting types of unseen primary and metastatic tumors with a single support sample from each class

We first investigated the impact of training sets with different primary types and different numbers of *N*-way predictions (*N* = 6, 8, and 10) on the prediction performance of primary cancers. The accuracy of 1-Nearest Neighbor (1-NN) was also computed, which is widely adopted benchmark model for one-shot and few-shot learning models. For an *N*-way prediction, 1-NN calculates the Euclidean distance between gene expressions of the query and a support sample and selects the type that has the minimum distance.

For all three CancerSiamese networks outperformed 1-NN for all three different *N*-way predictions (**Figure 3**A). Particularly, CancerSiamese-19 achieved the highest performance with 89.67%, 87.32%, and 84.59% accuracy for 6, 8, and 10-way predictions, respectively. They also represent 7%-8% improvement over 1-NN. Since the training and testing sets contain disjoint cancer types, these high accuracies suggest that there are discriminative gene expression markers shared among all cancer types that CancerSiamese models have successfully learned from training data, and predicting testing samples using these markers. In addition, we also observed that increasing diversity of training samples from 9 to 19 types improved accuracy, suggesting an improved learned representation of similar and dissimilar cancer types.

The test accuracies of CancerSiamese-MET were next compared to CancerSiamese-19 and 1-NN on metastatic cancer predictions (**Figure 3**B). We observe that CancerSiamese-MET achieved the best performance all different *N*-way predictions (~4-5% over 1-NN) but both CancerSiamese-MET and CancerSiamese-19 outperformed 1-NN (**Figure 4**B), demonstrating once again the ability of CancerSiamese in learning similarity/dissimilarity between cancer types. However, the accuracies are in the low 60%, indicating that expression signatures that define similar/dissimilar metastatic types learned from 10 training classes may not yet be fully generalized well to discriminate the signatures in testing samples. This could be partially due to the smaller metastatic training samples and also potentially higher expression heterogeneity in metastatic tumors, or lower sample diversity in training.

**Figure 4:**
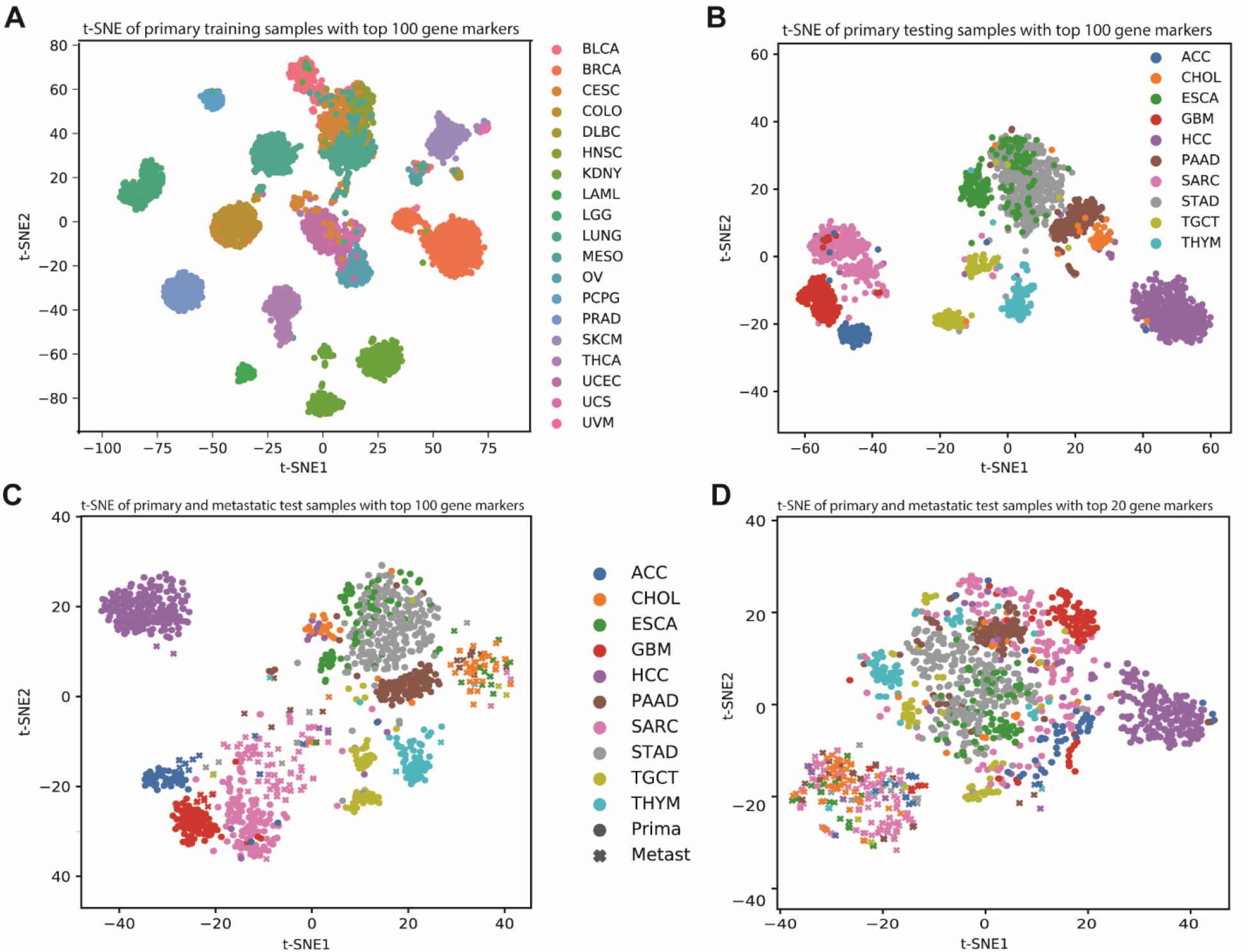
t-SNE plots of primary and metastatic test samples with top gene markers. A) primary training samples with top 100 gene markers. B) primary testing samples with top 100 gene markers. C) primary and metastatic test samples together with the top 100 gene markers. D) primary and metastatic test samples together with the top 20 gene markers.

Interestingly, CancerSiamese-19 (65.75%) reported only 2% reduction from the accurary of CancerSiamese-MET (63.46%), suggesting that the marker expression signatures learned from the 19 primary types shared significant similarity with those learned from the 10 metastatic types. In fact, the 10 metastatic types overlap with 10 of the 19 primary types. To further verify if the 10 overlapping types have shared expression signatures, we trained a 1D-CNN using the training samples from these 10 primary types and tested it on the training samples of the 10 metastatic types. As expected, this 1D-CNN was able to predict the metastatic tumors with an accuracy of 83.33%. The confusion matrix (Supplementary **Figure 3**) further delineates the shared gene expression signatures between primary and metastatic tumors. This result agreed with earlier studies that have also shown majority genes’ expression of primary and metastasized tumors resemble each other [23–25].

### 3.3 Identification and analysis of primary cancers marker genes learned by CancerSiamese

The test results have shown that CancerSiamese learned unique expression markers, which are shared among all cancer types between disjoint training and testing sets. Because CancerSiamese-19 had the best accuracy, we first selected it to investigate how CancerSiamese makes the prediction and to uncover the marker genes learned by CancerSiamese. To this end, we randomly selected up to 3,000 unique pairs of expressions from the same type for each of the 19 primary cancer types and used GBSM to calculate 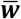 (See Methods), whose elements represent the scores of the corresponding genes to be marker genes (the ranked list of genes and scores can be found in Supplementary **Table S4**). To determine the set of marker genes, we performed a stepwise greedy forward selection (SGFS) [26], a popular feature selection method. Specifically, we ranked the genes in decreasing order based on their score. Next, we performed 1-NN for 6-way prediction on the test samples using multiple of 10 genes from the top of the list (**Table 2**). We selected the top 100 genes as the marker genes (Supplementary **Table S4**) as they gave the best accuracy (82.38%) is virtually the same as using all genes (82.47%, **Figure 3**A). t-SNE plots of training and testing samples using these marker genes further confirmed their discriminative power (**Figure 4**A&B) as samples from the same type are mostly grouped in clearly separated clusters. The heatmap of the markers demonstrated cancer-type specific expression patterns (Supplementary **Figure S2**). We then performed Gene Ontology (GO) and pathway enrichment analysis using DAVID, v6.7[27]. To ensure a meaningful functional annotation, we slightly relaxed the criterion and analyzed the top 5% (243 genes) of the ranked genes. As shown in Supplementary **Table S5**, the top functional annotation included apoptosis, cell growth, response to oxidative stress, extracellular matrix (ECM), and response to wounding. Apoptosis and cell growth are key determinants of cancer cell proliferation [28]. ECM and wound healing not only are critical component/indicator for the migration and metastasis of tumors [29], but also represent the fundamentally different tumor microenvironment between solid and hematopoietic cancers. The oxidative stress is known to trigger tumor progression and modulate the response to chemotherapies [30, 31]. Taken together, the top marker genes capture critical functions executed by tumors to survive and metastasize, thus potentially marking the differences between tumors and between primary and metastatic tumors.

### 3.4. Identification of metastatic tumor marker genes learned by CancerSiamese

Because CancerSiamese-19 achieved 63.46% accuracy for 6-way prediction for metastatic types, we first wondered if the primary markers might also be markers for metastatic cancer types. To this end, we performed 1-NN for 6-way prediction on the metastatic samples again using multiple of 10 genes in the ranked list of the primary cancers from the top but the accuracies stayed between 37-39% (**Table 2**), which is considerably lower than 63.46%. This result indicates that these primary markers are predominantly markers for primary but not metastatic cancer types. The t-SNE plot of the testing primary and metastatic samples (**Figure 4**C) further confirmed this finding. Indeed, many studies have also pointed out increased genome heterogeneity in metastasized tumors due to their different tumor microenvironment and immunological conditions [32, 33]. Interestingly, we observed that metastatic samples of 6 types (PAAD, STAD, ESCA, CHOL, SARC, and ACC) were separated from those of their corresponding primary cancers (**Figure 4**C). This further implies that these 100 primary marker genes also carry discriminate expression patterns between metastatic from primary samples, for at least these 6 cancer types. To validate this observation, we performed the SGFS on these 100 genes. For each increment of 10 genes in the ranked list, we trained a Gaussian Naïve Bayes (GNB) classifier to classify primary from metastatic cancer using the testing samples and determined that the top 20 genes were the discriminate markers (**Table 2**). The t-SNE plot of test samples using these 20 genes (**Figure 4**D) showed a clearer separation between primary and metastatic cancer samples than that using 100 markers (**Figure 4**C). We noted that many of the 20 genes were well-known to be strongly associated with metastatic mechanisms in many cancer types, such as *TAGLN2* [34], *S100A11* [35, 36], *CD74* [37], *TMSB10* [38, 39], and *ALDOA* [40].

To further determine the metastatic markers, we performed an SGFS on the CancerSiamese trained on the metastatic tumors with 10 classes and identified 250 marker genes (Supplementary **Table S4**), which produced an accuracy of 60.05% by 1-NN (Supplementary **Table S6**), which is very close to the accuracy by 1-NN from all genes (61.26%, **Figure 3**B). Comparing these metastatic marker genes with 100 markers for primary cancers yielded little overlapping (only 24 common genes), demonstrating the clear difference between metastatic and primary markers again. These different genes in the list of metastatic markers are responsible for the additional improvement over the 38.12% accuracy achieved by primary markers (**Table 2**). Functional annotation analysis of these 250 gene markers confirmed an association with cell-cell adhesion while also highlighted other fundamental molecular functions, such as nucleotide binding and peptide modification (Supplementary **Table S7**).

## 4. DISCUSSION AND CONCLUSION

In this paper, we proposed CancerSiamese, a one-shot learning model for predicting unseen primary and metastatic cancer type of a query expression sample. This model was developed based on the hypothesis that there exists a set of marker genes whose expressions define the similarity/dissimilarity between samples of the same/different types. The test results showed high prediction accuracies for primary cancers and 7-8% improvement over a 1-NN. These accuracies could be further improved by including more cancer classes in the training data. We determined 100 marker genes for the primary cancers and t-SNE plots further verified their cancer-type agnostic nature. Further analysis also showed that these maker genes are unique to the primary cancer types and functional enrichment analysis revealed that they are associated with many cancer-related functions.

In contrast, CancerSiamese-MET acquired accuracies only in low 60% for metastatic cancers, even though it still outperformed 1-NN by about 4%. Given that the number of MET500 samples for each cancer type is 10 to 20 fold less than those in TCGA (**Figure 1**), these low accuracies could very well be improved with an expanded collection of metastatic tumor samples and classes as witnessed in **Figure 3**A for the primary cancers. Nevertheless, metastatic tumor heterogeneity compounded by the impurity of metastatic tumor samples could further contribute to the low prediction accuracies. Despite the low performance, we still identified 200 marker genes, which account for 90% of the CancerSiamese’s performance. However, we observed little overlapping with the primary marker genes.

These results, especially those for primary cancers, serve to validate our hypothesis and, for the first time, demonstrated the possibility of applying one-shot learning for expression-based cancer type prediction. This new paradigm recognizes the reality of a small sample size in the era of precision oncology and provides a principled approach to meet the need for data-driven cancer diagnosis with small samples. Extension of CancerSiamese into a more versatile few-shot learning model will have a more direct impact on the practical application of CancerSiamese for precision tumor diagnosis. As one of the key machine learning challenges and opportunities for precision oncology and general precision medicine research lies in the small sample size, this work could inspire new and ingenious development of one-shot and few-shot learning solutions for improving cancer therapy and our understanding of cancer.

## References

1. Cancer Genome Atlas Research, N., et al., The Cancer Genome Atlas Pan-Cancer analysis project. Nat Genet, 2013. 45(10): p. 1113–20.

2. Robinson, D.R., et al., Integrative clinical genomics of metastatic cancer. Nature, 2017. 548(7667): p. 297–303.

3. Prasad, V., Perspective: The precision-oncology illusion. Nature, 2016. 537(7619): p. S63.

4. Joseph, M., M. Devaraj, and C.K. Leung. DeepGx: deep learning using gene expression for cancer classification. in 2019 IEEE/ACM International Conference on Advances in Social Networks Analysis and Mining (ASONAM). 2019. IEEE.

5. Lyu, B. and A. Haque. Deep learning based tumor type classification using gene expression data. in Proceedings of the 2018 ACM international conference on bioinformatics, computational biology, and health informatics. 2018.

6. Bazgir, O., et al. REFINED (REpresentation of Features as Images with NEighborhood Dependencies): A novel feature representation for Convolutional Neural Networks. arXiv e-prints, 2019. arXiv:1912.05687.

7. Fatima, N. and L. Rueda, iSOM-GSN: An Integrative Approach for Transforming Multi-omic Data into Gene Similarity Networks via Self-organizing Maps. Bioinformatics, 2020.

8. Sharma, A., et al., DeepInsight: A methodology to transform a non-image data to an image for convolution neural network architecture. Sci Rep, 2019. 9(1): p. 11399.

9. Mostavi, M., et al., Convolutional neural network models for cancer type prediction based on gene expression. BMC Med Genomics, 2020. 13(Suppl 5): p. 44.

10. Chiu, Y.C., et al., Deep learning of pharmacogenomics resources: moving towards precision oncology. Brief Bioinform, 2019.

11. Fei-Fei, L., R. Fergus, and P. Perona, One-shot learning of object categories. IEEE transactions on pattern analysis and machine intelligence, 2006. 28(4): p. 594–611.

12. Lake, B., et al. One shot learning of simple visual concepts. in Proceedings of the annual meeting of the cognitive science society. 2011.

13. Jeon, M., et al., ReSimNet: drug response similarity prediction using Siamese neural networks. Bioinformatics, 2019. 35(24): p. 5249–5256.

14. Koh, W. and S.J.b. Hoon, MapCell: Learning a comparative cell type distance metric with Siamese neural nets with applications towards cell-types identification across experimental datasets. 2019: p. 828699.

15. Nourani, E., et al., TripletProt: Deep Representation Learning of Proteins based on Siamese Networks. 2020.

16. Chung, Y.-A. and W.-H.J.a.p.a. Weng, Learning deep representations of medical images using siamese CNNs with application to content-based image retrieval. 2017.

17. Chen, M., et al., Multifaceted protein-protein interaction prediction based on Siamese residual RCNN. Bioinformatics, 2019. 35(14): p. i305–i314.

18. Zheng, W., et al., SENSE: Siamese neural network for sequence embedding and alignment-free comparison. Bioinformatics, 2019. 35(11): p. 1820–1828.

19. Ma, T. and A. Zhang. AffinityNet: semi-supervised few-shot learning for disease type prediction. in Proceedings of the AAAI Conference on Artificial Intelligence. 2019.

20. Koch, G., Richard Zemel, and Ruslan Salakhutdinov. Siamese neural networks for one-shot image recognition. in ICML deep learning workshop. 2015.

21. Springenberg, J.T., et al., Striving for simplicity: The all convolutional net. arXiv preprint arXiv: 1412.6806, 2014.

22. Chollet, F., keras. 2015.

23. Suzuki, M. and D. Tarin, Gene expression profiling of human lymph node metastases and matched primary breast carcinomas: clinical implications. Mol Oncol, 2007. 1(2): p. 172–80.

24. Iwamoto, T., et al., Distinct gene expression profiles between primary breast cancers and brain metastases from pair-matched samples. Sci Rep, 2019. 9(1): p. 13343.

25. Ho, T.H., et al., Differential gene expression profiling of matched primary renal cell carcinoma and metastases reveals upregulation of extracellular matrix genes. Annals of Oncology, 2017. 28(3): p. 604–610.

26. Chandrashekar, G., F.J.C. Sahin, and E. Engineering, A survey on feature selection methods. Computers & Electrical Engineering, 2014. 40(1): p. 16–28.

27. Huang da, W., B.T. Sherman, and R.A. Lempicki, Systematic and integrative analysis of large gene lists using DAVID bioinformatics resources. Nat Protoc, 2009. 4(1): p. 44–57.

28. Lowe, S.W. and A.W. Lin, Apoptosis in cancer. Carcinogenesis, 2000. 21(3): p. 485–95.

29. Gilkes, D.M., G.L. Semenza, and D. Wirtz, Hypoxia and the extracellular matrix: drivers of tumour metastasis. Nat Rev Cancer, 2014. 14(6): p. 430–9.

30. Saha, S.K., et al., Correlation between Oxidative Stress, Nutrition, and Cancer Initiation. Int J Mol Sci, 2017. 18(7).

31. Sosa, V., et al., Oxidative stress and cancer: an overview. Ageing Res Rev, 2013. 12(1): p. 376–90.

32. Voena, C. and R. Chiarle, Advances in cancer immunology and cancer immunotherapy. Discov Med, 2016. 21(114): p. 125–33.

33. Chitty, J.L., et al., Recent advances in understanding the complexities of metastasis. F1000Res, 2018. 7.

34. Han, M.Z., et al., TAGLN2 is a candidate prognostic biomarker promoting tumorigenesis in human gliomas. J Exp Clin Cancer Res, 2017. 36(1): p. 155.

35. Meding, S., et al., Tissue-based proteomics reveals FXYD3, S100A11 and GSTM3 as novel markers for regional lymph node metastasis in colon cancer. J Pathol, 2012. 228(4): p. 459–70.

36. Mori, M., et al., S100A11 gene identified by in-house cDNA microarray as an accurate predictor of lymph node metastases of gastric cancer. Oncol Rep, 2004. 11(6): p. 1287–93.

37. Greenwood, C., et al., Stat1 and CD74 overexpression is co-dependent and linked to increased invasion and lymph node metastasis in triple-negative breast cancer. J Proteomics, 2012. 75(10): p. 3031–40.

38. Zhang, X., et al., Thymosin beta 10 is a key regulator of tumorigenesis and metastasis and a novel serum marker in breast cancer. Breast Cancer Res, 2017. 19(1): p. 15.

39. Xiao, R., et al., TMSB10 promotes migration and invasion of cancer cells and is a novel prognostic marker for renal cell carcinoma. Int J Clin Exp Pathol, 2019. 12(1): p. 305–312.

40. Ji, S., et al., ALDOA functions as an oncogene in the highly metastatic pancreatic cancer. Cancer Lett, 2016. 374(1): p. 127–135.

